# THE CIRCADIAN CLOCK IN THE RETINAL PIGMENT EPITHELIUM CONTROLS THE DIURNAL RHYTHM OF PHAGOCYTIC ACTIVITY

**DOI:** 10.1101/2020.12.02.408799

**Authors:** Christopher DeVera, Jendayi Dixon, Micah A. Chrenek, Kenkichi Baba, P. Michael Iuvone, Gianluca Tosini

**Affiliations:** Department of Pharmacology & Toxicology and Neuroscience Institute, Morehouse School of Medicine, Atlanta, GA; Department of Ophthalmology and Emory Eye Center, Emory University, Atlanta, GA

**Keywords:** retinal pigment epithelium, retina, phagocytosis, aging, circadian clocks, Bmal1

## Abstract

The diurnal peak of phagocytosis by the retinal pigment epithelium (RPE) of photoreceptor outer segments (POS) is under circadian control, and it is believed that this process involves interactions from both the retina and RPE. Previous studies have demonstrated that a functional circadian clock exists within multiple retinal cell types and RPE cells. Thereby, the aim of the current study was to determine whether the circadian clock in the retina and or RPE controls the diurnal phagocytic peak of photoreceptor outer segments and whether selective disruption of the circadian clock in the RPE would affect RPE cells function and the viability during aging. To that aim, we first generated and validated an RPE tissue-specific KO of the essential clock gene*, Bmal1*, and then we determined the daily rhythm in phagocytic activity by the RPE in mice lacking a functional circadian clock in the retina or RPE. Then using electroretinography, spectral domain-optical coherence tomography, and optomotor response measurements of visual function we determined the effect of *Bmal1* removal in young (6-month old) and old (18-month old) mice. RPE morphology and lipofuscin accumulation was also determined in young and old mice. Our data show that the circadian clock in the RPE controls the daily diurnal phagocytic peak of POS. Surprisingly, the lack of a functional RPE circadian clock or the diurnal phagocytic peak does not result in any detectable age-related degenerative phenotype in the retina or RPE. Thus, our results demonstrate that the loss of the circadian clock in the RPE or the lack of the daily peak in phagocytosis of POS does not result in deterioration of photoreceptors or the RPE during aging.

## Introduction

Circadian clocks are present in almost all tissues and cells throughout the body (Takahashi, 2017), and the molecular mechanism responsible for the generation of circadian rhythms consists of two transcriptional translation feedback loops (TTFL) involving several genes (i.e., clock genes) and their protein products (Takahashi, 2017). Removal of clock gene *Bmal1* (i.e., *Arntl*) abolishes circadian rhythmicity at the behavioral and cellular level (Bunger et al., 2000) and induces premature aging, reduced life span, astrogliosis in the brain, and organ shrinkage (Kondratov et al., 2006; Musiek et al., 2013).

The mammalian eye also possesses a complete circadian system that controls many physiological functions within this organ (DeVera et al., 2019), and a few studies have also shown that *Bmal1* plays an important role in the maintenance of ocular health. Germline *Bmal1* KO mice show an increased incidence in cataract, cornea inflammation (Kondratov et al., 2006), and reduced photoreceptor viability during aging (Baba et al., 2018a). The circadian regulation of the retinal transcriptome is also dramatically affected by *Bmal1* removal since only a few genes – out of the thousand genes that are rhythmically expressed in wild-type mice – show some rhythmicity (Storch et al., 2007). Finally, in *Bmal1* KO mice the circadian regulation of the photic electroretinogram (ERG) is also absent (Storch et al., 2007). These same results are also obtained in mice lacking *Bmal1* only in the neural retina (Baba et al., 2018b; Storch et al., 2007). Retinal specific *Bmal1* KO mice show alterations in retinal circuitry and cone viability is significantly reduced during the aging process (Baba et al., 2018b). Additionally, a recent study has reported that *Bmal1* regulates spatial cone opsin expression in mouse retina through the activation of the gene encoding thyroid hormone deiodinase 2 (*Dio2*, Sawant et al., 2017) and regulates retinal neurogenesis (Sawant et al., 2019).

The retinal pigment epithelium (RPE) is constituted as a monolayer of post-mitotic epithelial cells that play an important role in the maintenance of photoreceptor health and functioning (Lakkaraju et al., 2020). Several studies have shown that the phagocytosis of the rod photoreceptor outer segment (POS) discs shows a daily peak occurring one to two hours after onset of light (Grace et al., 1999; LaVail, 1976; Lo and Bernstein, 1981), persists in constant darkness (Besharse et al., 1977; Grace et al., 1999; LaVail, 1976) and does not depend from the brain master circadian clock (i.e., the suprachiasmatic nuclei of the hypothalamus; Su Terman et al., 1993; Teirstein et al., 1980). Consistently with these earlier results, more recent studies have also demonstrated that RPE possesses a functional circadian clock (Baba et al., 2010, 2017). A previous study has also reported that in vivo removal of the mouse daily peak in phagocytic activity has a negative consequence on the health of the RPE and photoreceptors during aging (Nandrot et al., 2004), although more recent work suggests that the lack of the peak may not be detrimental for the health of the RPE and photoreceptors (Goyal et al., 2020a; Nandrot and Finnemann, 2008). In our previous study, we also reported that although the daily rhythm in phagocytic activity in the RPE of D_2_R KO mice is no longer present, clock genes expression in the RPE is not affected by the removal of D_2_R signaling (Goyal et al., 2020a). Therefore, we inferred that the lack of a negative phenotype was probably due to persistent circadian regulation of RPE function.

The aim of the current study was to first determine whether the circadian clock located in the RPE drives the diurnal peak in the phagocytosis of POS by RPE and then to determine whether disruption of the circadian clock in the RPE would negatively affect the health of RPE and of the photoreceptors during aging.

## Materials and Methods

### Animals

RPE^cre^ (rtTA-VMD2/Cre) (Le et al., 2008) and *Bmal1*^*fl/fl*^ (Jackson laboratory, 007668) were bred to generate the conditional and inducible Cre recombinase expression in retinal pigment epithelium tissue: RPE^cre^; *Bmal1*^*fl/fl*^. The following primer pairs were used to genotype for both the Cre locus (Forward: 5’- CATCGCTCGACCAGTTTAGTT-3’, Reverse: 5’- CTGACGGTGGGAGAATGTTAAT-3’) and the *Bmal1*^*fl/fl*^. locus (Forward: 5’-TCCTGGTTGGTCCAAGAATATG-3’, Reverse: 5’- CTGACCAACTTGCTAACAATTA-3’). RPE^cre^; *Bmal1*^*fl/fl*^. mice were fed either regular mouse lab diet (RPE *Bmal1* WT; Bioserv, S4207) or doxycycline (dox; RPE *Bmal1* KO; dox; Bioserv, S3888) mouse lab diet at p60-p74. Details about the production, maintenance, and genotyping of Chx10^cre^; *Bmal1*^*fl/fl*^ (Retina *Bmal1* KO) and floxed control (Retinal *Bmal1* WT; *Bmal1*^*fl/fl*^; Jackson Laboratory, 007668) are reported in our previous study (Baba et al., 2018b). All mice were housed on a 12:12 light:dark cycle with lights on at 6 AM [zeitgeber time [ZT]0) and lights off at 6 PM (ZT12). Additionally, RPE Bmal1 WT and RPE Bmal1 KO mice were aged to 6-months (young) and 18-months (old) of age. All animals were housed in the animal facility at Emory University School of Medicine. All experimental procedures were performed in accordance with the NIH Guide on Care and Use of Laboratory Animals and were approved by the Emory University and the Morehouse School of Medicine Animal Care and Use Committees.

### Phagosome counting assay

Whole eyes were processed for phagosome counting as previously described (Goyal et al., 2020a) and labeled with the primary rhodopsin 4D2 antibody (Abcam, ab98887). Briefly, whole eyes were fixed in 4% paraformaldehyde for at least 3 hours and transferred to 30% sucrose (Fisher, S5-500) for cryoprotection. Once sufficiently cryoprotected, eyes were embedded (Fisher, 4585), sectioned at 12 μm, and stored at −20 °C until processed. On the day of processing, goat serum (Vector, S-1000) and fragmented goat anti-mouse antibodies (Jackson Immuno Research, 115-007-003) was used for antigen blocking before overnight incubation at 4 °C with rhodopsin 4D2 (1:500, Abcam, ab98887). Slides were washed and incubated with goat anti-mouse 488 (1:1000, CST, 4408S) for up to 2 hours at room temperature (RT). Slides were washed and counterstained with DAPI (1:500, ThermoFisher, D1306). Immunopositive rhodopsin phagosomes localized to the RPE layer (confirmed with DAPI) were counted in 2×150 microns of the retina in the nasal to a temporal orientation on at least 4 eye sections with a confocal microscope (Zeiss, LSM 700).

### Western blot

RPE was isolated as previously described (DeVera and Tosini, 2020) then lysed in ice-cold RIPA buffer (Boston Bioproducts, BP-116X) with protease inhibitors. RPE protein homogenates (at least 10 μg/well) were separated on a tris/glycine/SDS 7.5% gradient mini-protean gel (Bio-Rad, 456-1026) and electrophoretically transferred to a nitrocellulose membrane (Bio-Rad, 1704156) using the Transblot turbo system (Bio-Rad, 1704150) for immunoblotting. Membranes were blocked for 1 hour at RT on a shaker in 5% BSA (Tocris, 5217) with 0.1% Tween-20 (Bio-Rad, 170-6531). Each membrane was incubated with primary antibodies (D_2_R, Millipore Sigma, AB5084P and Bmal1, Cell Signaling Technology, 14020S) in 5% BSA with 0.1% Tween-20 overnight at 4°C on a shaker. The membranes were washed three times for five minutes in 0.1 M TBS and 0.1% Tween-20 on a shaker at RT. A secondary antibody conjugated with HRP (anti-rabbit; CST, 7074S) were incubated at RT on a shaker for 1 hour. The membranes will be washed four times for five minutes in 0.1 M TBS and 0.1% Tween-20 on a shaker at RT. The development of the membranes will be accomplished with ECL western blot substrate (ThermoFisher Scientific, 32106) for five minutes at RT. Protein bands were visualized using image J (1.51w).

### Spectral domain-optical coherence tomography (SD-OCT) and fundus imaging

Retinal thickness measurements were performed as previously described (Baba et al., 2018b). Briefly, mice were anesthetized and a circular scan 0.57 mm from the optic nerve head was made (Phoenix Micron IV). Total retinal and photoreceptor receptor layer thicknesses were measured and quantified using Adobe Photoshop. Fundus imaging of mice was done at 6-month and 18-month of age using a confocal laser scanning ophthalmoscope (cLSO; Heidelberg Spectralis, 0226-HRAOCT2-MC) using the manufacturer provided BluePeak™ autofluorescence imaging module set at max laser intensity.

### Electroretinography (ERG)

Photic and scotopic ERG measurements were performed, as previously described in Baba et al., (2018b). Briefly, all mice were measured midday and anesthetized with ketamine/xylazine. Each recording epoch was 250 ms and each stimulus flash presented in an UTAS BigShot ganzfield (LKC Technologies). For dark-adapted conditions, a series of 5 stimulating flashes that ranged from 0.00039 to 25.3 cd*s/m^2^ with each flash lasting 20 μs. For light-adapted ERG recordings, a steady background-adapting field of 29.76 cd*s/m^2^ was presented to saturate rod photoreceptors. After 10 minutes of light-adaptation, each step consisted of 60 flashes of 3.14 cd*s/m^2^ that lasts 2 minutes for a total measurement time of 35 minutes. The amplitude of each a-wave was measured from baseline to trough and the b-wave from the trough to the peak of the response.

### Opto-motor response (OMR)

Mice underwent visual psychophysical testing using the OptoMotry device (CerebralMechanics, Inc., D430) to assess spatial frequency threshold and contrast sensitivity as previously described (Baba et al., 2018b).

### RPE flat mount morphology and autofluorescence analysis

RPE flat mount analysis was performed as previously described (Goyal et al., 2020a, 2020b). Briefly, each eye following enucleation was placed in Z-fix (Anatech, 170) for 10 minutes at room temperature and then rinsed up to five times with 0.01M phosphate-buffered saline. On a microscope slide (VWR, 16004-406), radial cuts around the limbus of the eye were made using spring scissors (WPI, 501235) to remove the anterior segment of the eye (cornea and lens). The remaining eyecup was divided into 4 petals and allowed to be flattened against the microscope slide. The now exposed retina was peeled by using Dumont #5/45 forceps (FST, 11251-35) and RPE flat mounts were placed in a 24 well plate (Fisher, 12565501) with 0.01M phosphate-buffered saline and processed immediately for zonula occludins-1 (ZO-1; 1:500, Millipore Sigma, MABT11) and Bmal1 (Cell Signaling Technologies, 14020S) immunostaining. RPE flat mounts were blocked and then incubated with ZO-1 antibody (1:500, Millipore Sigma, MABT11) overnight at 4°C. Flat mounts were washed and incubated in Alexa Fluor 488 (1:500, goat anti-rat, Invitrogen, A110006) for up to 2 hours on a shaker at room temperature. RPE cell junctions were visualized on a confocal microscope (Zeiss LSM 700) in the 488 nm channel, and autofluorescent particles were visualized in the 568 nm channel. In order to determine the number of autofluorescent particles per RPE cell, a pipeline in Cell Profiler (v 2.2.0) was created to: 1) identify RPE cells on the isolated RPE flat mount via ZO-1 staining, 2) identify all autofluorescent particles present on the RPE flat mount, 3) create a mask from the ZO-1 labeling to identify RPE cells, and 4) count the number of autofluorescent particles in each RPE cell. Additionally, RPE cell morphology parameters such as area, eccentricity, solidity, and compactness were measured from step 1 on isolated RPE cells, as previously described (Boatright et al., 2015). For all RPE morphology analyses, only central measurements were made on each RPE petal (up to 1.0 mm from the optic nerve head).

### Statistical analyses

For all datasets analyzed, tests for assumptions of normality by Shapiro-Wilk and assumptions of equal variance by the F-test was done prior to any statistical test is applied to assess for differences. Statistical analyses were done with either unpaired t-tests, one-way ANOVA, or two-way ANOVA. Tukey posthoc multiple comparisons test was done with significance set at p < 0.05 for all statistically significant ANOVA results with power > 0.8. Unpaired t-tests for comparison between two groups was set at p < 0.05 with power >0.8. All data were expressed as mean ± SEM (standard error of the mean).

## Results

### Removal of Bmal1 from RPE cells

RPE^cre^; Bmal1^fl/fl^ mice were feed with dox diet (i.e., RPE Bmal1 KO) or control diet (i.e., RPE Bmal1 WT) from post-natal day 60 to 74 (P60-P74; Figure 1A). Flat mount from an RPE Bmal1 WT mouse crossed with an Ai6(RLC-ZsGreen) cre reporter mouse model demonstrated about 95% cre recombinase expression (Figure 1B) with a majority of expression in RPE cells with some cre recombinase expression in endothelial cells as demonstrated in a transverse eye section (Figure 1C). BMAL1 immunoreactivity was abolished in the majority of the RPE cells (Figure 1E) and quantification of BMAL1 by western blot indicates that BMAL1 levels were reduced by more than 70% in RPE Bmal1 KO when compared to RPE Bmal1 WT mice (Figure 1F-G, unpaired t-test, p < 0.05).

**Figure 1.**
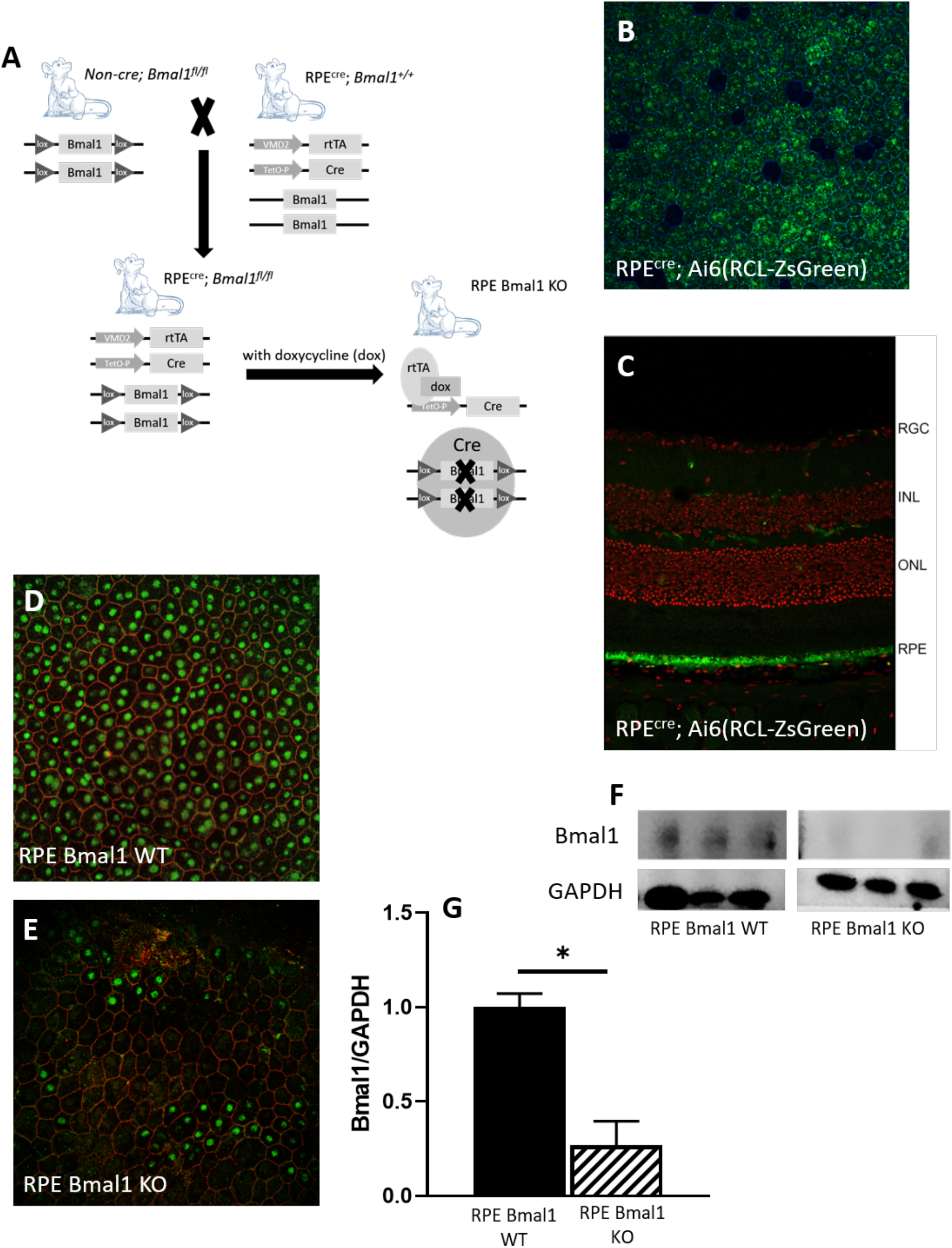
Generation and validation of RPE *Bmal1* KO mouse model. A) Graphical representation of the breeding schema for the generation of RPE^cre^; *Bmal1*^*fl/fl*^. mice and doxycycline (dox) treatment. B) We first confirmed cre recombinase activity by crossing RPE^cre^ mice and Ai6(RCL-ZsGreen) mice (Jackson Laboratory, cat no. 007906). When RPE cells were visualized as a flat mount *en face* view, there was about a 95.8% expression of cre recombinase (green) in RPE cells that were demarcated with zonula occludens (blue). C) When whole eyes from RPE^cre^; Ai6(RCL-ZsGreen) were prepared as transverse sections, an expected expression of cre recombinase (green) can be seen in the RPE cell layer with some unexpected cre recombinase expression in what is probably endothelial cells in the outer and inner plexiform layer as identified between the nuclear layers in the retina via propidium iodide staining (red). D-E) Visualization of representative microphotographs displaying Bmal1 immunoreactivity in RPE *Bmal1* KO and RPE *Bmal1* WT mice as prepared in an RPE flat mount. Bmal1 immunoreactivity in RPE cells was decreased (about 70%) as visualized on an RPE flat mount when compared to 100% Bmal1 expression in RPE Bmal1 WT mice. F) Representative Western blot bands for Bmal1 and Gapdh antibodies and densitometry analysis of the bands. G) Bmal1 protein was significantly reduced (about 70%) in RPE *Bmal1* KO when compared to RPE *Bmal1* WT mice (D; * = p < 0.05, unpaired t-test, n=3 mice/group). Bars on graphs represent mean ± SEM.

### Loss of Bmal1 in the RPE dramatically reduces the daily peak of phagocytic activity by the RPE

As shown in Figure 2, removal of Bmal1 from the retina did not affect the daily pattern of RPE phagocytic activity (Figure 2E), whereas removal of Bmal1 dramatically blunted the daily activity in phagocytic activity of RPE Bmal1 KO mice (Figure 2C). Interestingly, there were no differences in the total number of phagosomes engulfed in all mouse models used (Figure 2F, one-way ANOVA, p > 0.1). We have previously reported that activation of dopamine 2 receptor (D_2_R) signaling is required for the presence of the diurnal phagocytic peak of photoreceptor outer segments (Goyal et al., 2020a). Thus, we decided to investigate whether removal of Bmal1 from the RPE will also affect D_2_R expression. Consistent with our previous study, D_2_R expression was significantly reduced (about 70%) in RPE Bmal1 KO mice when compared to RPE Bmal1 WT mice when measured at ZT1 (Figure 3A-B; p < 0.05, unpaired t-test).

**Figure 2.**
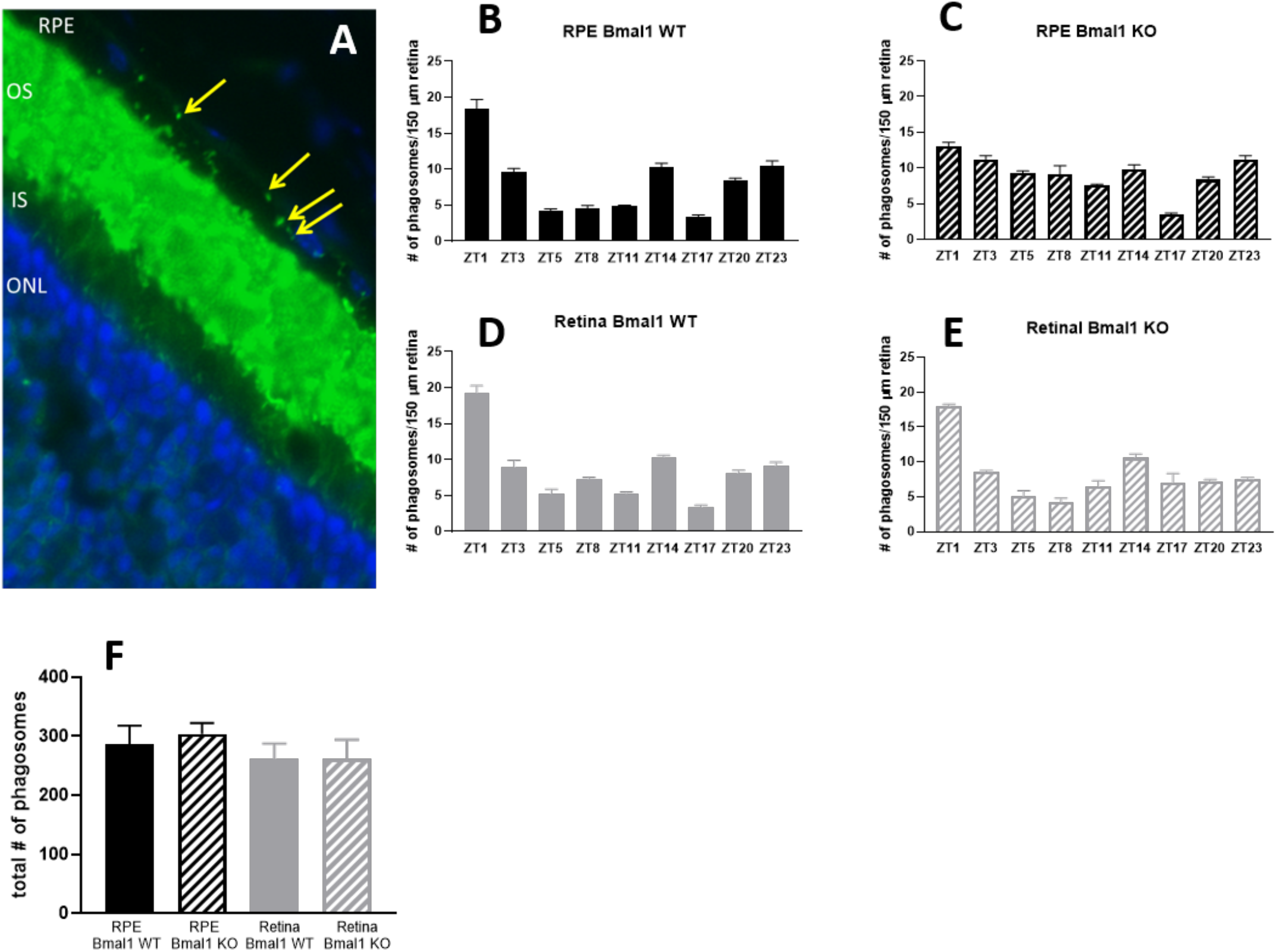
The RPE circadian clock controls the diurnal peak in phagocytosis of POS. A) Representative microphotograph of a transverse eye section immunostained with anti-rhodopsin antibody (Rho4D2, green) and DAPI (blue). Phagosomes (marked with yellow arrows) can be seen as small green particles present in the RPE cell layer. B-E) Bar graphs indicating the number of phagosomes per 150 μm of RPE at different time points in four different genotypes. A significant difference in the number of phagosomes at the different times of the day was observed in all the four genotypes (one-way ANOVA, Tukey post hoc, p < 0.05, but the amplitude of the diurnal peak (i.e., ZT1) in the number of phagosomes in RPE Bmal1 KO mice was dramatically dampened (C). No difference was observed in the total number of phagosomes engulfed during the day among the different genotypes (F; p > 0.1, one-way ANOVA). Bars represent the mean ± SEM (n = 4–6/group). OS = outer segments; IS = inner segments; ONL = outer nuclear layer.

**Figure 3.**
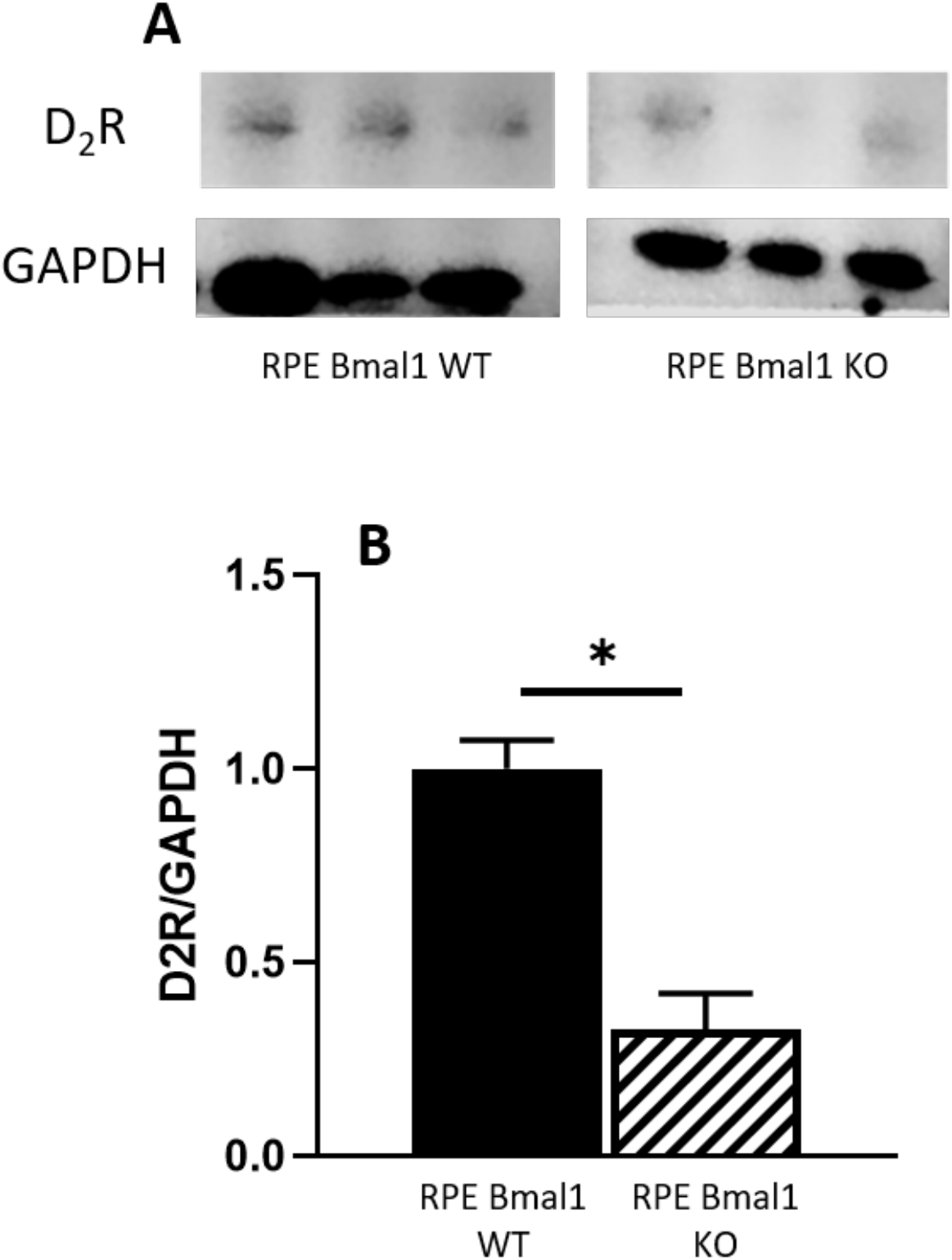
Dopamine 2 receptor protein is decreased in RPE *Bmal1* KO mice. A) Representative Western blot bands for D_2_R and Gapdh antibodies; B) Densitometry analysis of the band intensities indicated a significant reduction in D_2_R protein in RPE *Bmal1* KO mice when compared to RPE *Bmal1* WT mice (* = p < 0.05, unpaired t-test, n=3/group). Bars on graphs represent mean ± SEM.

### Removal of Bmal1 from the RPE does not affect retinal structure during aging

Total retina thickness did not differ between the young RPE Bmal1 KO compared to RPE Bmal1 WT (two-way ANOVA, p > 0.1, Figure 4B) while old RPE Bmal1 KO shows a slight increase in total retina thickness (two-way ANOVA, Tukey posthoc, p < 0.05, Figure 4B). No significant differences were observed in the thickness of the photoreceptors layer between the two genotypes and ages (Figure 4C; p > 0.1, two-way ANOVA). Analysis of the scotopic and photopic ERGs did not indicate any significant differences in the amplitudes of the a- and b-waves between the two genotypes at both ages (two-way ANOVA, p > 0.1 in all cases, Figures 5A-C). No difference among the two genotypes was also observed in the visual acuity and contrast sensitivity at both ages (two-way ANOVA, p > 0.1 in all cases; Figures 6A-B).

**Figure 4.**
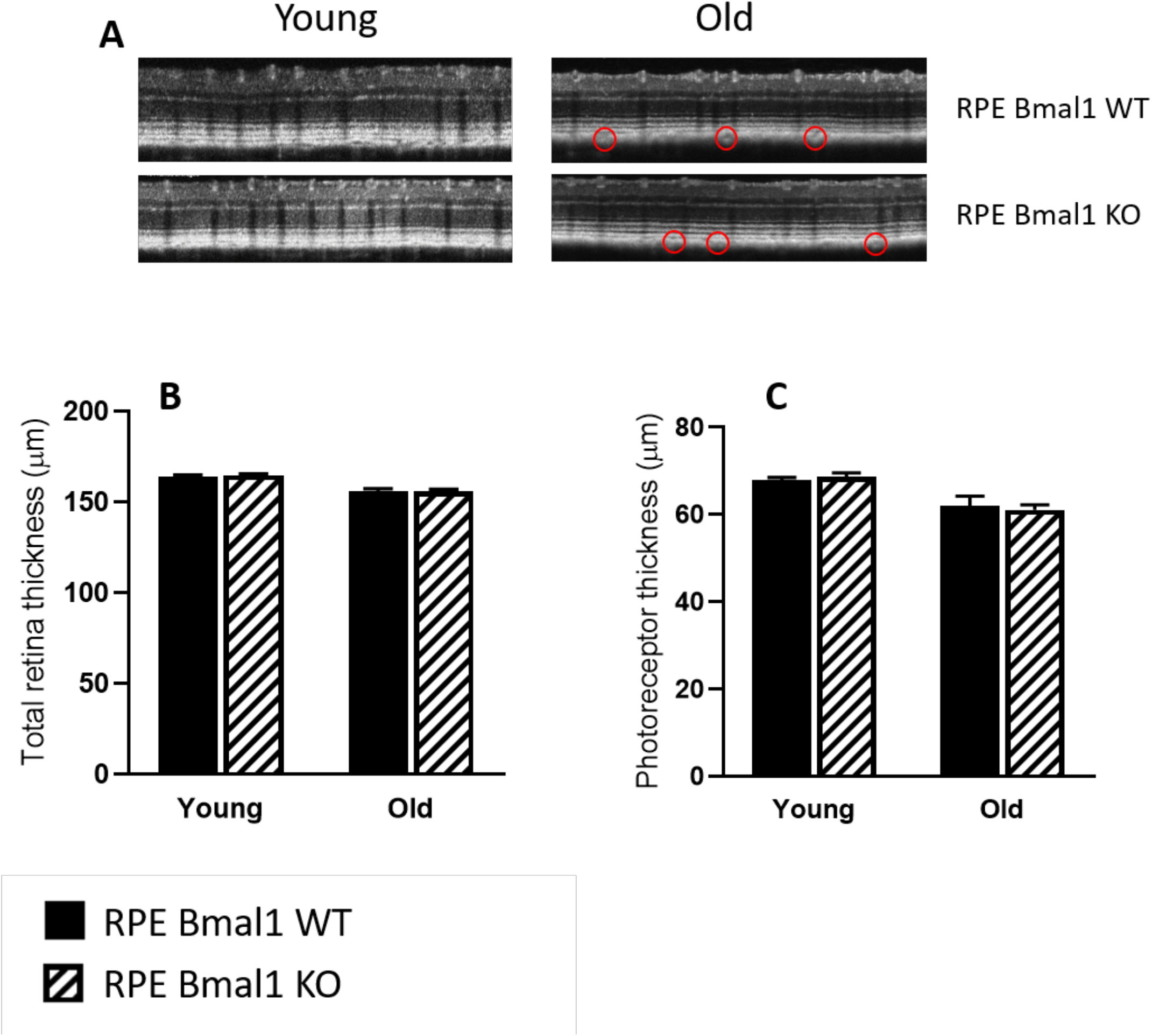
Retinal thickness is not affected by disruption of the RPE circadian clock. A) Representative retinal images taken with SD-OCT from young and old RPE *Bmal1* WT and RPE *Bmal1* KO mice. B) Total retinal thickness was not affected in the RPE of mice lacking *Bmal1* in the RPE at any age measured (p > 0.1, two-way ANOVA, Tukey posthoc, n=4-6/group). C) No differences were observed in the thickness of the photoreceptor layer between the two genotypes at both ages (p > 0.1, two-way ANOVA. n=4-6 mice/group). Small pockets can be observed in the layer between the basal layer of the RPE and the apical choroid in both aged groups but absent in the younger group of mice (red circles). Bars on graphs represent mean ± SEM.

**Figure 5.**
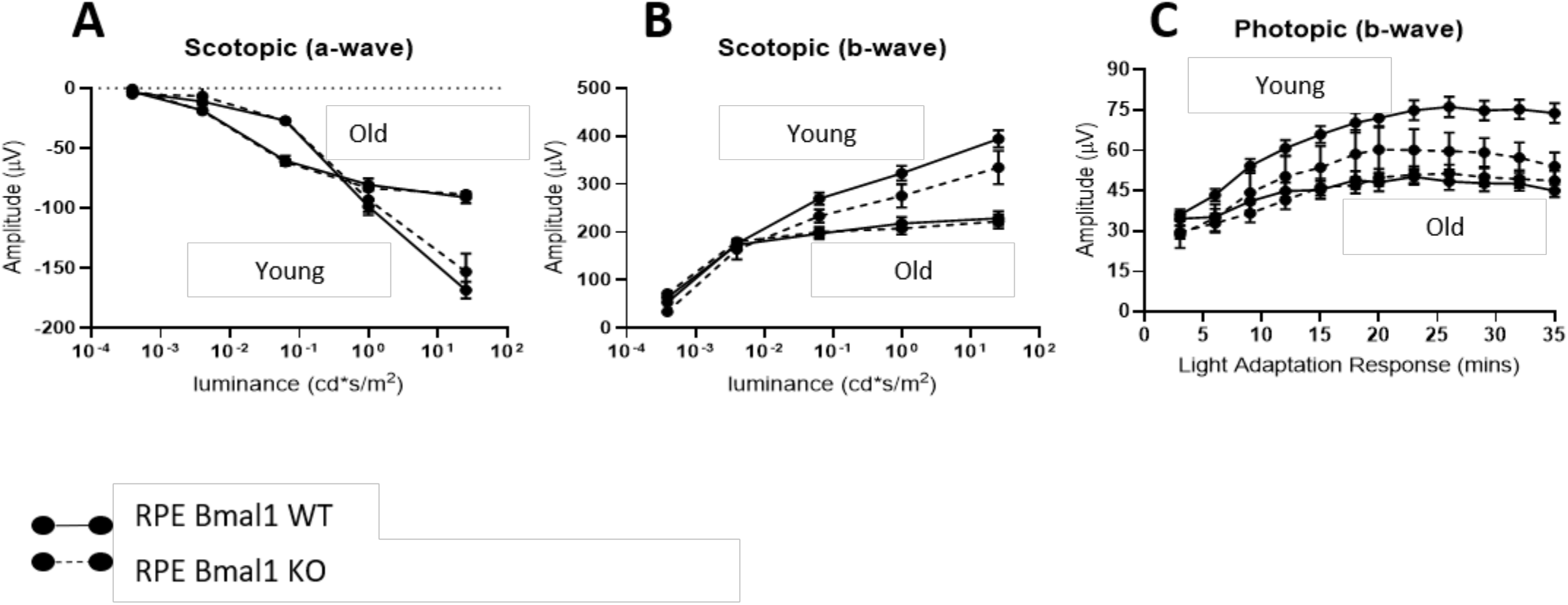
Functioning of rods and cones are not affected by the loss of *Bmal1* in the RPE. Retina function was assessed with electroretinography at the level of both rod (i.e., scotopic) and cone (i.e., photopic) photoreceptors. A-B) At all ages considered, there was no difference in rod photoreceptor function in RPE *Bmal1* KO when compared to RPE *Bmal1* WT (p > 0.1, two-way ANOVA, n=4-6/group). C) Additionally, cone photoreceptor function was also no affected in RPE *Bmal1* KO when compared to RPE Bmal1 WT mice at all ages (p > 0.1, two-way ANOVA). Circle on graphs represent mean ± SEM.

**Figure 6.**
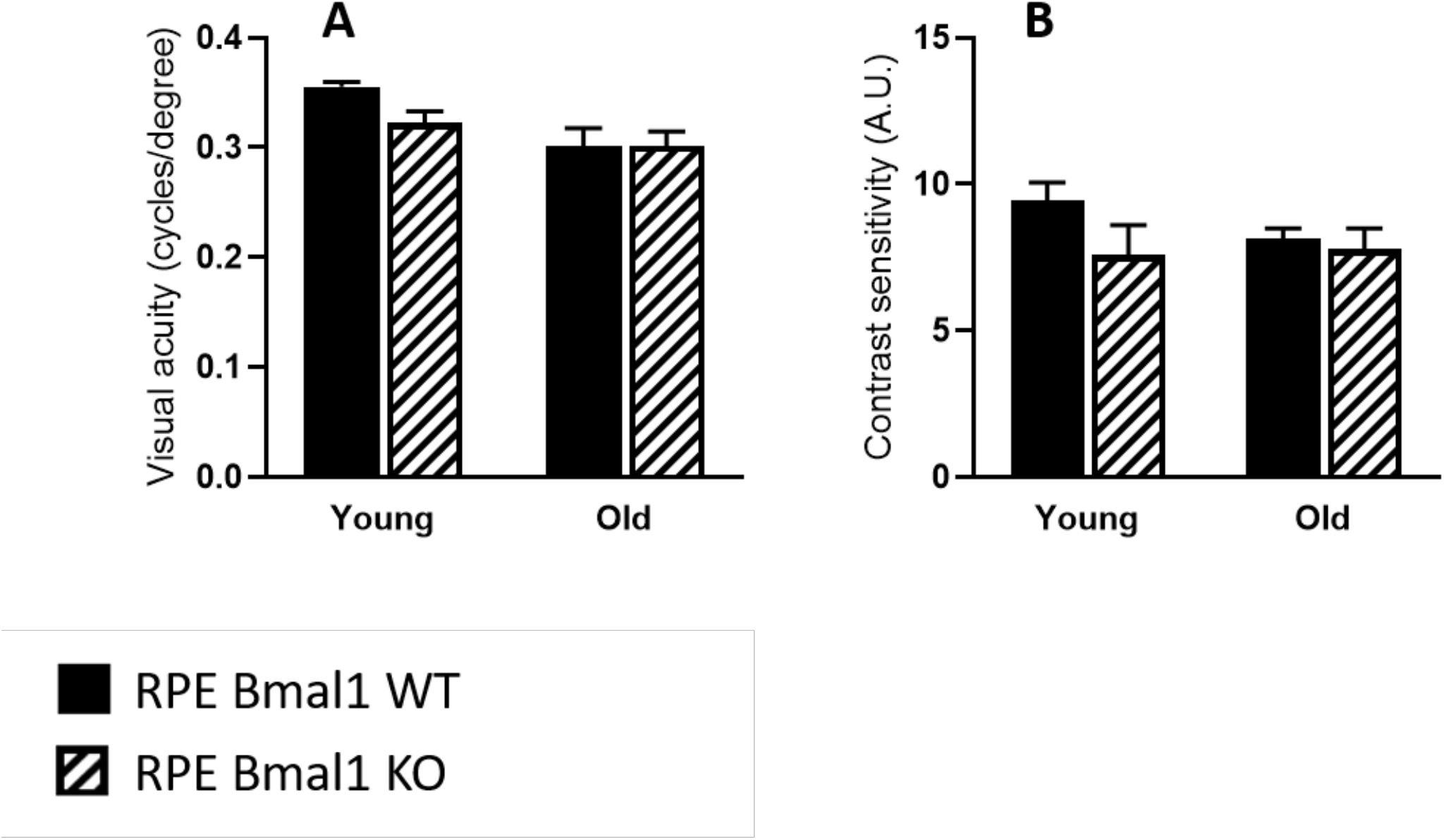
Psychophysical of visual function is not affected in the absence of *Bmal1* in RPE cells. A) No differences were observed in visual acuity (A, p > 0.1, two-way ANOVA) in RPE *Bmal1* KO mice when compared to similarly aged RPE *Bmal1* WT mice. B) Contrast sensitivity was also not affected in RPE *Bmal1* KO when compared to RPE *Bmal1* WT at all the ages tested (p > 0.1, two-way ANOVA, n=4-6/group). Bars on graphs represent mean ± SEM.

### Removal of Bmal1 does not produce morphological abnormalities in the RPE

Analysis of autofluorescent particles in the apical part of the RPE revealed no significant differences between the RPE Bmal1 KO and RPE Bmal1 WT at both ages (Figure 7F, unpaired t-test, p > 0.1). Finally, no significant changes in RPE morphological parameters were observed in old RPE Bmal1 KO with respect to age-matched RPE Bmal1 WT mice (unpaired t-tests, p > 0.1 in all cases; Figures 8A-D).

**Figure 7.**
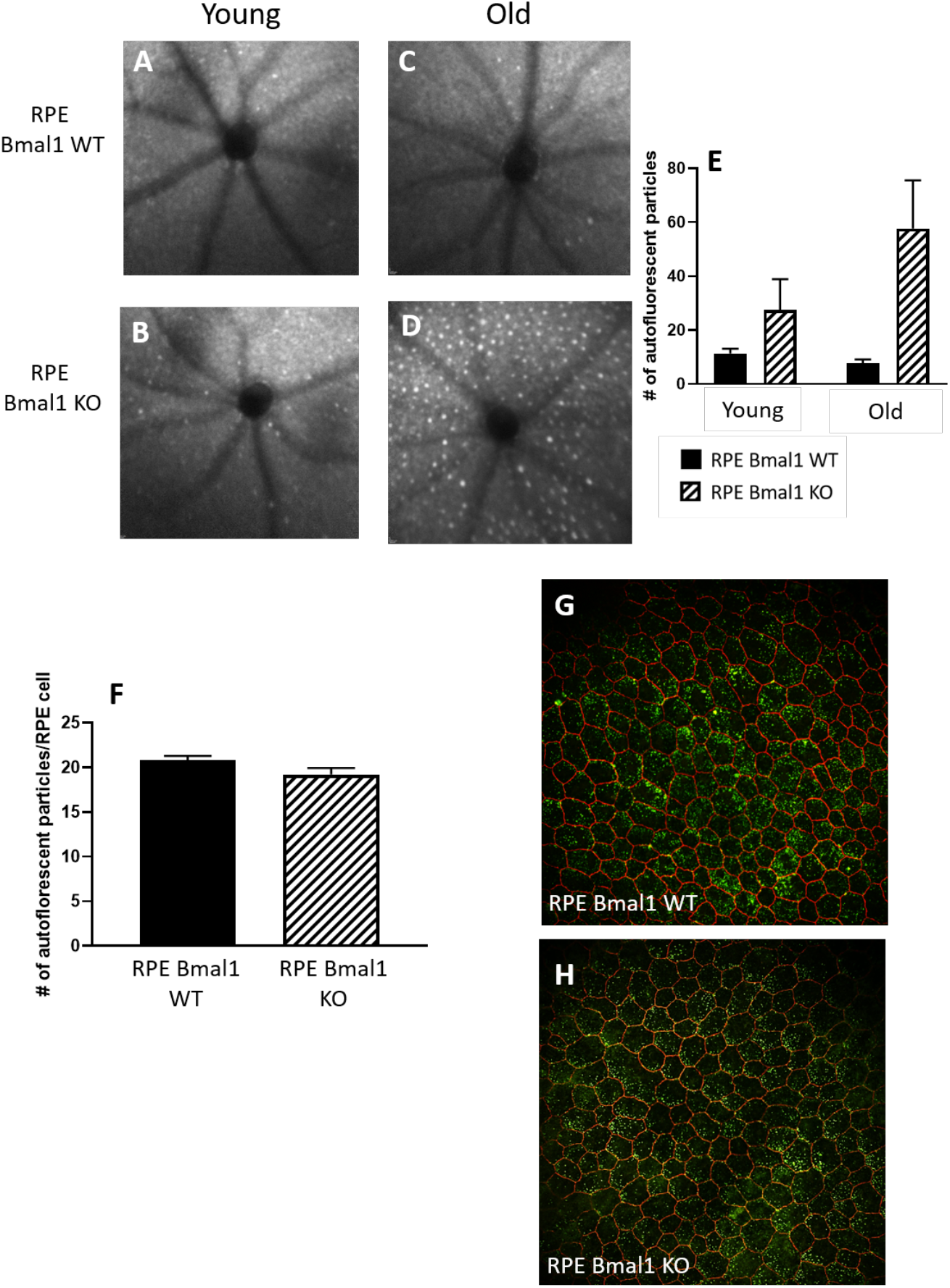
Autofluorescence particle accumulation in the fundus or RPE cells is not affected by disruption of the RPE circadian clock. A-D) The health of the mouse eye fundus was visualized with a confocal laser scanning ophthalmoscope in RPE *Bmal1* KO and RPE *Bmal1* WT in both young and old mice. E) There was no significant difference in the number of bright white autofluorescent particles when RPE *Bmal1* KO mice compared to RPE *Bmal1* WT mice at all ages (p > 0.1, two-way ANOVA). G-H) We also quantified the number of apical autofluorescent particles in isolated RPE cells in both old RPE Bmal1 WT mice and RPE Bmal1 KO mice. F) No differences were observed in the number of accumulated apical autofluorescent particles as a result of the loss of the RPE circadian clock (p > 0.1, unpaired t-test, n=5-6/group). Bars on graphs represent mean ± SEM.

**Figure 8.**
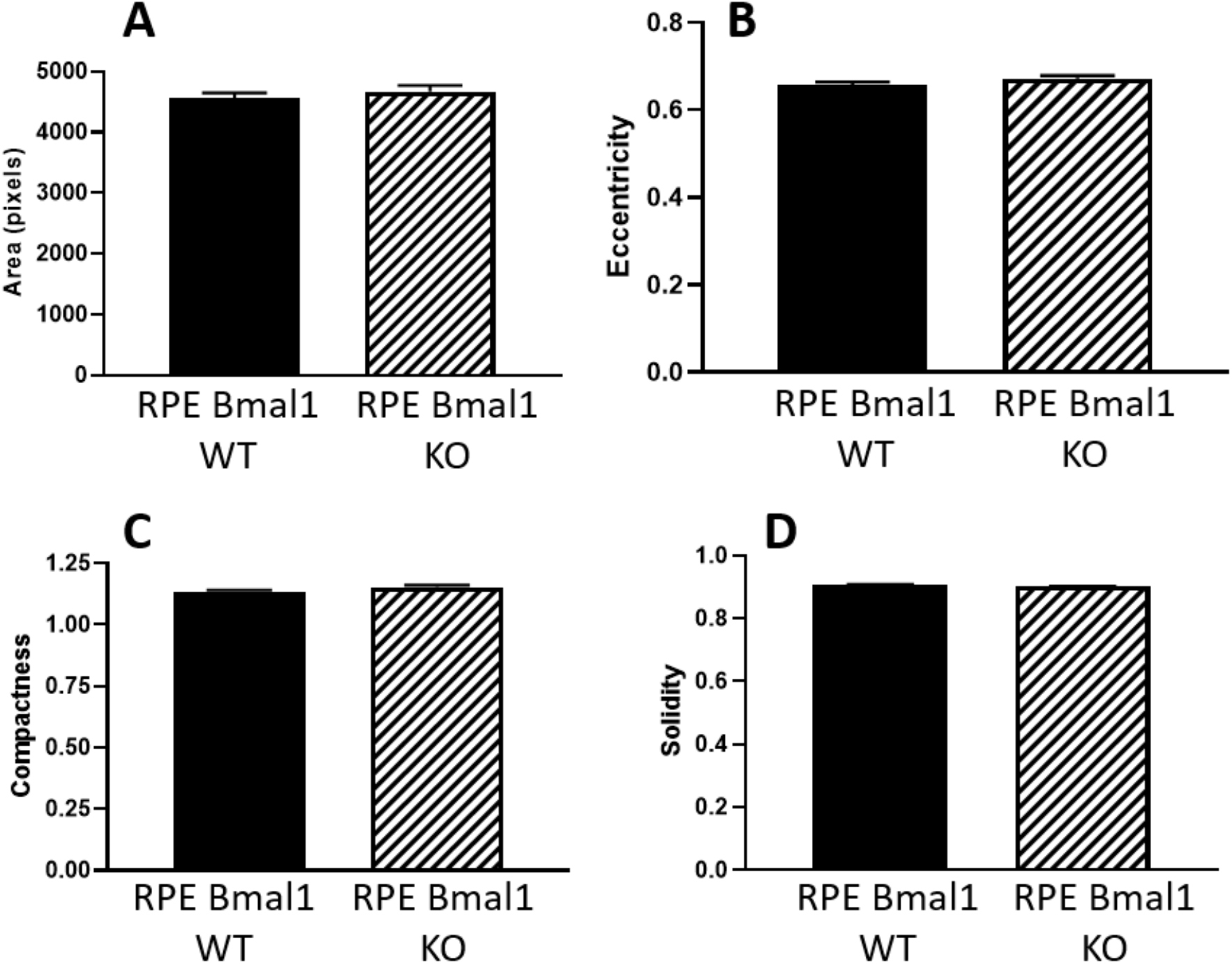
RPE morphology is not affected with loss of *Bmal1* in the RPE. A-D) Using cell profiler (v2.2.0), we were able to measure the area, eccentricity, compactness, and solidity of RPE cells. No differences were observed in any of the four shape parameters measured in old RPE *Bmal1* KO mice compared to RPE *Bmal1* WT mice (p > 0.05, unpaired t-test, n=5-6/group). Bars on graphs represent mean ± SEM.

## Discussion

The circadian clock system in the eye plays an important role in the modulation of several important ocular functions (DeVera et al., 2019), and the daily and/or circadian regulation of the phagocytic activity of the POS by RPE is probably the most studied of these rhythms (Besharse and Hollyfield, 1979; Goyal et al., 2020a; Grace et al., 1999; Laurent et al., 2017; LaVail, 1976; Nandrot et al., 2004; Terman et al., 1993; Teirstein et al., 1980). However, it is still unclear whether the presence of the phagocytic peak 1-2 hours after the onset of light is important for the health of the RPE and photoreceptors and whether this rhythm is controlled by the retinal or RPE circadian clock or whether the presence of both clocks is necessary. The findings of the present study indicate that a functional circadian clock in the RPE is sufficient and necessary for the presence of the daily peak in phagocytic activity by the RPE (Figure 2), whereas disruption of the retinal circadian clock does not affect this rhythm. Our data also confirm our previous study (Goyal et al., 2020a) by showing that the loss of the diurnal phagocytic peak does not produce any detrimental phenotype in the retina or RPE. Surprisingly, our study also indicated that the lack of a functional circadian clock in the RPE does not produce any detectible negative consequence on the RPE even 16 months after the circadian clock has been disrupted in these cells. Interestingly, although cre recombinase expression was very high (96%), the reduction of *Bmal1* protein levels was lower (about 70%) and not homogenous since BMAL1 immunoreactivity in the RPE cells of the KO mice showed a mosaic pattern (Figure 1). These data are consistent with a previously published study (Le et al., 2008).

A recent study has reported the presence of a second peak in the phagocytic activity soon after the onset of darkness (around ZT14, Lewis et al., 2018) and, interestingly, the presence of this “nocturnal peak” was still present in RPE and retina *Bmal1* KO mice (Figure 2). While the biological significance of this second peak in the nocturnal period is still undetermined, it is noteworthy to mention that our recent work has shown that this peak is not present in D_2_R KOs mice, but no deleterious phenotypes in either the retina or RPE were observed (Goyal et al., 2020a). Finally, our data provide additional support for a key role of dopamine via D_2_R signaling in the regulation of the morning peak in phagocytic activity by RPE cells since the loss of *Bmal1* in RPE cells results in diminished levels of D_2_R protein (Figure 3) and thus providing a potential mechanism to explain the loss of the diurnal phagocytic peak of POS in the RPE *Bmal1* KO mice.

Many studies have shown that 10-20% of the mouse transcriptome is under circadian clock control in many tissues/organs (Hughes et al., 2017), including the RPE (DeVera and Tosini, 2020; Louer et al., 2020a). Not surprisingly, the transcriptional processes under circadian regulation in the RPE include many of the pathways known to be involved in the regulation of phagocytic activity and metabolism, such as actin cytoskeleton remodeling (Bulloj et al., 2013; Louer et al., 2020a), integrin signaling (Goyal et al., 2020a; Nandrot et al., 2004), cAMP signaling (Mustafi et al., 2013), protein phosphorylation (Lakkaraju et al., 2020) and mitochondrial electron transport chain (DeVera and Tosini, 2020; Louer et al., 2020b). Hence, it is quite surprising that the removal of a functional circadian clock from the RPE did not induce any negative phenotype. This lack of a negative phenotype is particularly surprising given the nature of RPE cells (post-mitotic), the high metabolic rate and the functions that these cells play in the regulation of many biological processes (e.g., recycling of visual photopigments, phagocytosis of POS, homeostasis of the interphotoreceptor matrix, etc.).

Although the number of autofluorescent particles in RPE cells or the fundus of the mice – which includes the layers of retina, outer retina, and optic disc – was not different between the two genotypes at both ages (Figure 7), we did notice that the old mice in both genotypes displayed small empty pockets that can be observed between the RPE cells and choroid layers in the SD-OCT images (red circles; Figure 4A). The biological significance of these small accumulations of pockets on the basal side of RPE cells is still being investigated, but it is worth noting that these small pockets can be observed in aged human eyes and speculated to be a biomarker for early age-related macular degeneration (Heesterbeek et al., 2020). Additionally, our study also demonstrates that while autofluorescence might be observed in the fundus in the posterior area of the eyes, it does not necessarily translate to functional deficits.

As we have previously mentioned, removal of *Bmal1* negatively affects the function and viability of many cell types in the brain and the eye (Baba et al., 2018a; 2018b; Kondratov et al., 2006; Musiek et al., 2013; Sawant et al., 2017, 2019; Storch et al., 2007) and therefore our new results are somewhat puzzling. This unexpected result can be explained by several previous studies i) post-embryonic removal of *Bmal1* may not produce negative results as observed in germline *Bmal1* removal (Yang et al., 2016); *ii)* rhythmic metabolic processes may persist in a tissue even in the absence of *Bmal1* (Ray et al., 2020); *iii)* the presence of *Bmal1* in 30% of RPE cells may still drive rhythmic gene expression in the entire tissue (Figure 1), and finally *iv)* the unchallenging housing condition (e.g., light, food, etc.,) at which the mice are maintained may mitigate the consequence of a dysfunctional circadian clock (Turek et al., 2005). Further studies will be needed to address this important point and thus to fully understand the role played by the circadian clock in the regulation of RPE functions.

In conclusion, our study using a tissue-specific *Bmal1* KO mouse model demonstrates that the circadian clock in the RPE controls the daily diurnal peak in phagocytosis of POS. In addition, our data also indicate that loss of the diurnal phagocytic peak of POS or a functional RPE circadian clock does not result in any deleterious effects in the retina and RPE during aging. Finally, our study also suggests that the disruption of the circadian clock in a tissue-specific may not always produce expected negative consequences from the use of circadian disruption.

## Notes

### Competing Interest Statement

The authors have declared no competing interest.

